# Transcriptional profiling of the *Candida auris* response to exogenous farnesol exposure

**DOI:** 10.1101/2021.08.23.457447

**Authors:** Ágnes Jakab, Noémi Balla, Ágota Ragyák, Fruzsina Nagy, Fruzsina Kovács, Zsófi Sajtos, Andrew M. Borman, István Pócsi, Edina Baranyai, László Majoros, Renátó Kovács

**Affiliations:** Department of Molecular Biotechnology and Microbiology, Institute of Biotechnology, Faculty of Science and Technology, University of Debrecen, Debrecen, Hungary; Department of Medical Microbiology, Faculty of Medicine, University of Debrecen, Debrecen, Hungary; Doctoral School of Pharmaceutical Sciences, University of Debrecen, Debrecen, Hungary; Department of Inorganic and Analytical Chemistry, Agilent Atomic Spectroscopy Partner Laboratory, University of Debrecen, Debrecen, Hungary; UK National Mycology Reference Laboratory, Public Health England, Science Quarter, Southmead Hospital, Bristol BS10 5NB, UK; Medical Research Council Centre for Medical Mycology (MRC CMM), University of Exeter, Exeter EX4 4QD, UK; Faculty of Pharmacy, University of Debrecen, Debrecen, Hungary

**Keywords:** *Candida auris*, farnesol, quorum-sensing, transcriptome analysis, oxidative stress, metal, iron, zinc, copper

## Abstract

The antifungal resistance threat posed by *Candida auris* necessitates bold and innovative therapeutic options. Farnesol, a quorum-sensing molecule with a potential antifungal and/or adjuvant effect; it may be a promising candidate in alternative treatment regimens. To gain further insights into the farnesol-related effect on *C. auris*, genome-wide gene expression analysis was performed using RNA-Seq. Farnesol exposure resulted in 1,766 differentially expressed genes. Of these, 447 and 304 genes with at least 1.5-fold increase or decrease in expression, respectively, were selected for further investigation. Genes involved in morphogenesis, biofilm events (maturation and dispersion), gluconeogenesis, iron metabolism, and regulation of RNA biosynthesis showed down-regulation, whereas those related to antioxidative defense, transmembrane transport, glyoxylate cycle, fatty acid β-oxidation, and peroxisome processes were up-regulated. In addition, farnesol treatment increased the expression of certain efflux pump genes, including *MDR1*, *CDR1*, and *CDR2*. Growth, measured by change in CFU number, was significantly inhibited within 2 hours of the addition of farnesol (5.8×10^7^±1.1×10^7^ and 1.1×10^7^±0.3×10^7^ CFU/ml for untreated control and farnesol-exposed cells, respectively) (*p*<0.001). In addition, farnesol treatment caused a significant reduction in intracellular iron (152.2±21.1 vs. 116.0±10.0 mg/kg), manganese (67.9±5.1 vs. 18.6±1.8 mg/kg), and zinc (787.8±22.2 vs. 245.8±34.4 mg/kg) (*p*<0.05–0.001) compared to untreated control cells, whereas the level of cooper was significantly increased (274.6±15.7 vs. 828.8±106.4 mg/kg) (*p*<0.001). Our data demonstrate that farnesol significantly influences the growth, intracellular metal ion contents, and gene expression related to fatty acid metabolism, which could open new directions in developing alternative therapies against *C. auris*.

**Importance:** *Candida auris* is a dangerous fungal pathogen that causes outbreaks in health care facilities, with infections associated with high mortality rate. As conventional antifungal drugs have limited effects against the majority of clinical isolates, new and innovative therapies are urgently needed. Farnesol is a key regulator molecule of fungal morphogenesis, inducing phenotypic adaptations and influencing biofilm formation as well as virulence. Alongside these physiological modulations, it has a potent antifungal effect alone or in combination with traditional antifungals, especially at supraphysiological concentrations. However, our knowledge about the mechanisms underlying this antifungal effect against *C. auris* is limited. This study has demonstrated that farnesol enhances the oxidative stress and reduces the fungal survival strategies. Furthermore, it inhibits manganese, zinc transport, and iron metabolism as well as increases fungal intracellular copper content. In addition, metabolism was modulated towards β-oxidation. These results provide definitive explanations for the observed antifungal effects.

## Introduction

A dramatic increase in resistance to conventional antifungal agents has been reported for *Candida auris* worldwide, leading to evasion from efficient therapeutic options. The current COVID-19 pandemic situation may further promote the spreading of this fungal superbug. Superinfections by *C. auris* in critically ill COVID-19 patients have been related to high 30-day mortality rates, usually above 50% (1–3).

Farnesol is a fungal quorum-sensing molecule inducing hyphae-yeast morphological switching in *C. albicans* (4). In the past decade, several studies have reported that farnesol can generate oxidative stress and influence membrane permeability and cellular polarization in certain fungal species, especially at supraphysiological concentrations (5–7). Although farnesol does not affect the growth rate of *C. albicans* when growing in planktonic form, it significantly decreased the growth of *C. auris* regarding both planktonic cells and also 1-day-old biofilms of this organism (7). Recently, alternative therapeutic approaches designed to disturb quorum-sensing have become an attractive treatment strategy (8–9). The usage of farnesol and traditional antifungal drugs in combination may provide new insights into the management of newly emerged fungal species, such as *C. auris*, which poses a global threat to the nosocomial environment (7,10).

Previously, our group has reported the potential therapeutic benefit of farnesol against *C. auris* (7,10). However, there are no data to date that describe the total transcriptome changes induced by farnesol. Such data might help elucidate the *C. auris*-specific response to exogenous farnesol exposure. To gain further insights into previously described physiological consequences of farnesol treatment, we determined genome-wide gene expression changes induced by farnesol exposure using total transcriptome sequencing (RNA-Seq).

## Materials and methods

### Fungal strain, media, and culture conditions

The *C. auris* isolate 12 (NCPF 8973), belonging to the South Asian/Indian lineage, was obtained from the National Mycology Reference Laboratory (United Kingdom) (11). The test strain was maintained and cultured on yeast extract peptone dextrose (YPD) agar [1% yeast extract (Alfa Aesar, USA), 2% mycological peptone (Oxoid, UK), 2% dextrose (VWR International Llc., Hungary) and ± 2% agar (VWR International Llc., Hungary), pH 5.6] as described previously (12).

To study the effect of farnesol on short-term transcriptional response, *C. auris* pre-cultures were grown in 5 mL YPD medium at 30°C at a shaking frequency of 2.3 Hz for 18 hours. Subsequently, the inoculum was diluted to an optical density of 0.1 at λ = 640 nm (OD_640_) with YPD (at 0 hours incubation time as defined in growth assays), and the cultures were further grown at 37°C and 2.3 Hz shaking frequency. At 4 hours incubation time, the cultures were supplemented with 75 μM farnesol, and microbial growth was monitored by measuring changes in OD_640_ and colony-forming unit (CFU) (7,13). Farnesol (Merck, Budapest, Hungary) was obtained as 3M stock solution that was diluted to a 30 mM working stock solution in 100% methanol. The working concentrations were prepared in YPD. Farnesol-free control flasks contained 1% (v/v) methanol. Growth was evaluated in six independent experiments and is presented as the mean ± standard deviation (SD). Statistical comparison of growth-related data was performed by paired Student’s *t* test. The differences between values for treated and control cells were considered significant at a *p* value < 0.05.

### RNA isolation and sequencing

Total RNA was extracted from untreated control cells and 75 μM farnesol-treated cultures in three biological replicates. Briefly, fungal cells were collected at 2 hours following farnesol exposure by centrifugation (5 min at a relative centrifugal force [RCF] of 4,000 × g at 4°C). The cells were washed three times with phosphate-buffered saline (PBS) and stored at −70°C until use. Total RNA samples were prepared from freeze-dried cells (CHRIST Alpha 1‐2 LDplus lyophilizer, Osterode, Germany) derived from untreated and farnesol-treated cultures using TRISOL (Invitrogen, Austria) reagent, according to Chomczynski et al. (1993) (14). To determine the final RNA concentration and quality, samples were analyzed on an Agilent BioAnalyzer using the Eukaryotic Total RNA Nano Kit (Agilent Technologies, Inc., Santa Clara, CA, USA) according to the manufacturer’s protocol. Samples with RNA integrity number (RIN) values > 7 were accepted for the library preparation process. Three independent cultures were used for RNAseq experiments and RT-qPCR tests.

To obtain global transcriptome data, high-throughput mRNA sequencing was performed. The RNA-Seq libraries were prepared from total RNA using the NEBNext Ultra II RNA Sample preparation kit (NEB, USA) according to the manufacturer’s protocol. The single-read 75-bp-long sequencing reads were generated on an Illumina NextSeq500 instrument. Approximately 18–22 million reads per samples were generated. The library preparations and the sequencing run were performed by the Genomic Medicine and Bioinformatics Core Facility of the Department of Biochemistry and Molecular Biology, Faculty of Medicine, University of Debrecen, Hungary. Raw reads were aligned to the reference genome (genome: „http://fungi.ensembl.org/candida_auris_gca_002759435/Info/Index”;features:”http://www.candidagenome.org/download/gff/C_auris_B8441/archive/C_auris_B8441_version_s01-m01-r11_features_with_chromosome_sequences.gff.gz”), and aligned reads varied between 90 and 95% in each sample. The DESeq algorithm (StrandNGS software) was used to obtain normalized gene expression values. Gene expression differences between farnesol-exposed and control groups were compared by a moderated *t* test; the Benjamini-Hochberg false discovery rate was used for multiple-testing correction, and a corrected *p* value of < 0.05 was considered significant (differentially expressed genes). Up- and down-regulated genes were defined as differentially expressed genes with > 1.5-fold change (FC, up-regulated genes) or < −1.5-FC (down-regulated genes) values. The FC ratios were calculated from the normalized gene expression values.

### Reverse transcriptase-quantitative polymerase chain reaction (RT-qPCR) assays

Changes in the transcription of selected oxidative stress response, membrane transport, virulence, and primary metabolism genes were validated by RT-qPCR (13). The RT-qPCRs with Luna Universal One-Step RT-qPCR kit (NEB, USA) were performed according to the protocol of the manufacturer, using 500 ng of DNase- (Sigma, Budapest, Hungary) treated total RNA per reaction. Oligonucleotide primers (Table S1) were designed with the software packages Oligo Explorer (version 1.1.) and Oligo Analyzer (version 1.0.2). Three parallel measurements were performed with each sample in a LightCycler® 96 real-time PCR instrument (Roche, Switzerland). Relative transcription levels (ΔΔCP value) were calculated as ΔCPcontrol − ΔCPtreated, where ΔCPcontrol = CPtested gene, control − CPreference gene, control for untreated control, and ΔCP_treated_ = CP_tested gene, treated_ − CP_reference gene, treated_ for farnesol-exposed cultures (13). The CP values represent the RT-qPCR cycle numbers of crossing points. The reference gene used was *ACT1* (B9J08_000486). The ΔΔCP values are expressed as mean ± SD calculated from three independent measurements, and ΔΔCP values significantly (*p* < 0.05) higher or lower than zero were determined using the Student *t*-test.

### Functional enrichment analysis

Gene set enrichment analyses on the up-regulated and down-regulated gene sets were performed with *Candida* Genome Database Gene Ontology Term Finder (https://www.candidagenome.org/cgi-bin/GO/goTermFinder), using function, process, and component gene ontology (GO) terms. Only hits with a *p* value of < 0.05 were considered in the evaluation process (Table S2).

Besides GO terms, groups of functionally related genes were also generated by extracting data from the *Candida* Genome Database (https://www.candidagenome.org) unless otherwise indicated. The enrichment of *C. auris* genes from these gene groups in the up-regulated and down-regulated gene sets was tested with Fisher’s exact test (*p* < 0.05). The following gene groups were created:

i. Virulence-related genes – Genes involved in the genetic control of *C. albicans* virulence were collected according to Mayer et al. (2013), Höfs et al. (2016), and Araújo et al. (2017) (15–17).
ii. Metabolic pathway-related genes – This group contains all genes related to the carbohydrate, ergosterol, and fatty acid biochemical pathways according to the pathway databases (https://pathway.candidagenome.org/).
iii. Antioxidant enzyme genes – This group includes genes encoding functionally verified and/or putative antioxidant enzymes according to catalases (GOID:4096), SODs (GOID:4784), glutaredoxins (GOID:6749), thioredoxins (GOIDs:8379 and 51920), and peroxidases (GOID:4601) GO terms.
iv. Iron metabolism-related genes – Genes involved in iron acquisition by *C. albicans* were collected according to Fourie et al. (2018) (18).
v. Zinc, manganese, and copper homeostasis genes – Genes involved in zinc and copper acquisition were collected according to Gerwien et al. (2018) (19).

The complete gene lists of the above-mentioned gene groups are available in Supplementary Table 3.

### Assays of iron, manganese, zinc, and copper contents of *C. auris* cells

*C. auris* pre-cultures were grown, and farnesol exposure was performed as described above. Yeast cells were collected by centrifugation (5 min, 4,000 × g, 4°C) after 2 hours of incubation following farnesol exposure. Changes in fungal dry cell mass (DCM) were determined after freeze-drying (20). The metal contents of the dry biomass were measured by inductively coupled plasma optical emission spectrometry (ICP-OES; 5110 Agilent Technologies, Santa Clara, CA, USA) following atmospheric wet digestion in 3 mL of 65% (M/M) HNO_3_ and 1 mL of 30% (M/M) H_2_O_2_ in glass beakers. The metal contents of the samples were calculated and expressed in DCM units (mg/kg), as described previously by Jakab et al. 2021 (20). The metal contents of the biomasses were determined in triplicate, and mean ± SD values were calculated. Statistical significance of changes was determined by two-way ANOVA. Significance was defined as a *p* value of < 0.05.

## Results

### Growth-related experiments

The growth of *C. auris* was examined following 75 μM farnesol treatment in YPD. Adding farnesol to pre-incubated cells resulted in a remarkable growth inhibition, starting at 6 hours post inoculation, which was confirmed by both absorbance (OD_640_) measurements and CFU determination. Growth was significantly inhibited within 2 hours of the addition of farnesol as assessed both by CFU changes (5.8 × 10^7^ ± 1.1 × 10^7^ and 1.1 × 10^7^ ± 0.3 × 10^7^ CFU/ml for untreated control and farnesol-exposed cells, respectively) (*p* < 0.001) and observed absorbance values (1.28 ± 0.04 and 0.72 ± 0.04 for untreated control and farnesol-exposed cells, respectively, at OD 640 nm) (*p* < 0.001) (Fig. 1). The observed growth inhibition was further confirmed by changes in measured DCM at 12 hours incubation time (5.5 ± 0.2 and 1.3 ± 0.1 g/L for untreated control and farnesol-exposed cells, respectively) (*p* < 0.001).

**Figure 1.**
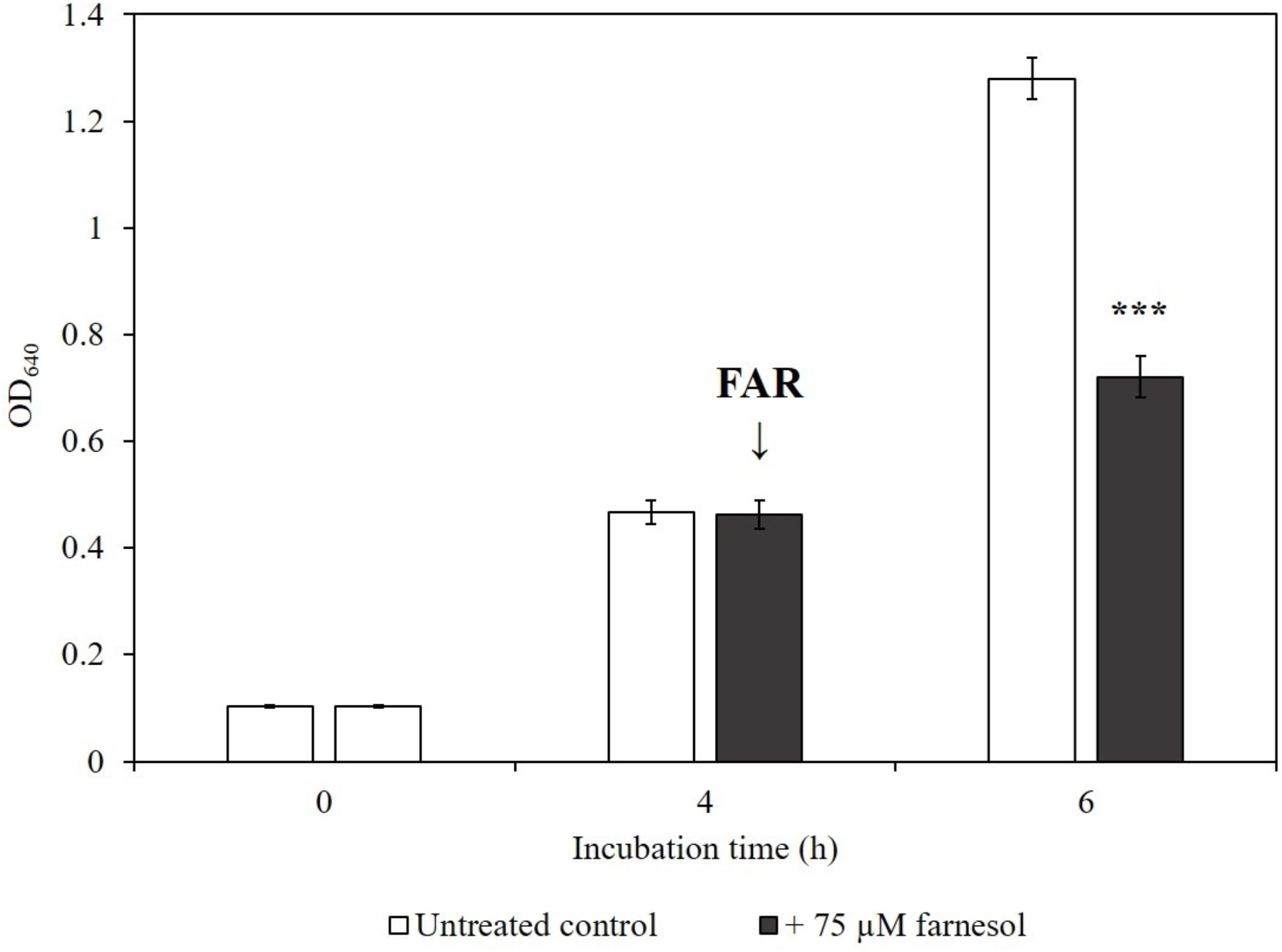
Effect of farnesol on the growth of *C. auris*. Changes in the growth of *C. auris* were monitored by measurement of the absorbance (OD_640_). Following a 4 h incubation time, farnesol was added at 75 μM final concentration to the YPD cultures. Data represent mean values ±SD calculated from 6 independent experiments. The asterisks indicate a statistically significant difference between control and farnesol-treated cultures calculated by paired Student’s *t* test.

### Transcriptional profiling and RNA-Seq data validation

Principal component analysis (PCA) and hierarchical clustering were performed to provide a visual representation of the transcriptomic similarities between samples treated with farnesol and the untreated controls (Figs. 2A and B). Samples from different conditions (with or without farnesol) clustered separately, whereas those from the same conditions clustered together, indicating a high level of correlation among samples as well as distinctive transcriptome profiles. Analyses of the RNA sequencing data clearly indicated that farnesol has a remarkable effect on *C. auris* gene expression, leading to significant alterations in the transcriptome.

**Figure 2.**
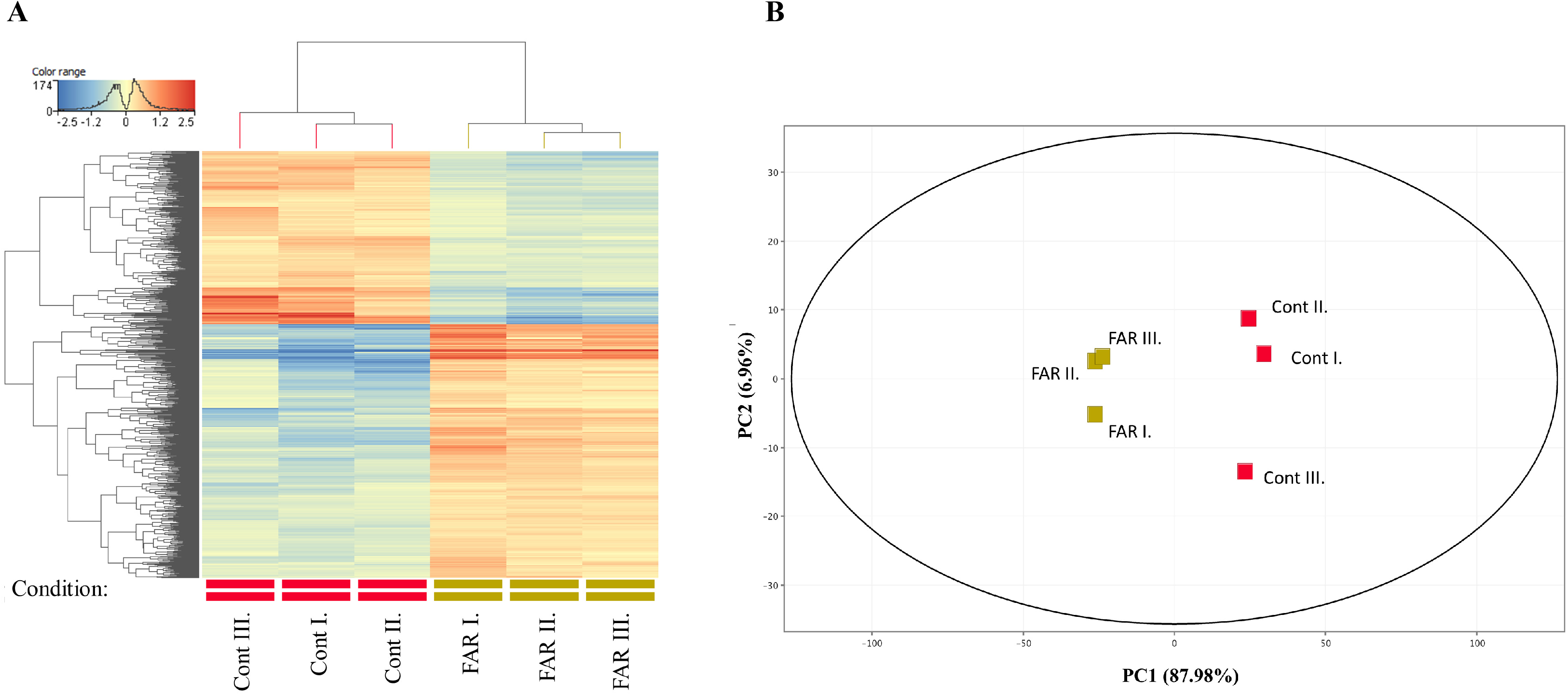
Cluster (A) and principal component (B) analysis of the transcriptome data. Symbols represent untreated control (Cont) and 75 μM farnesol exposure (FAR) cultures. The distribution of transcriptome data obtained in three independent series of experiments (I, II and III). Analyses were performed with the StrandNGS software using default settings.

Comparison of the farnesol-exposed *C. auris* global gene expression profile with that of unexposed cells revealed 1,766 differentially expressed genes. Among those, 447 were up-regulated and 304 were down-regulated in the farnesol-exposed samples compared to the untreated controls (Figs. 3 and 4, Tables S2 and 3).

**Figure 3.**
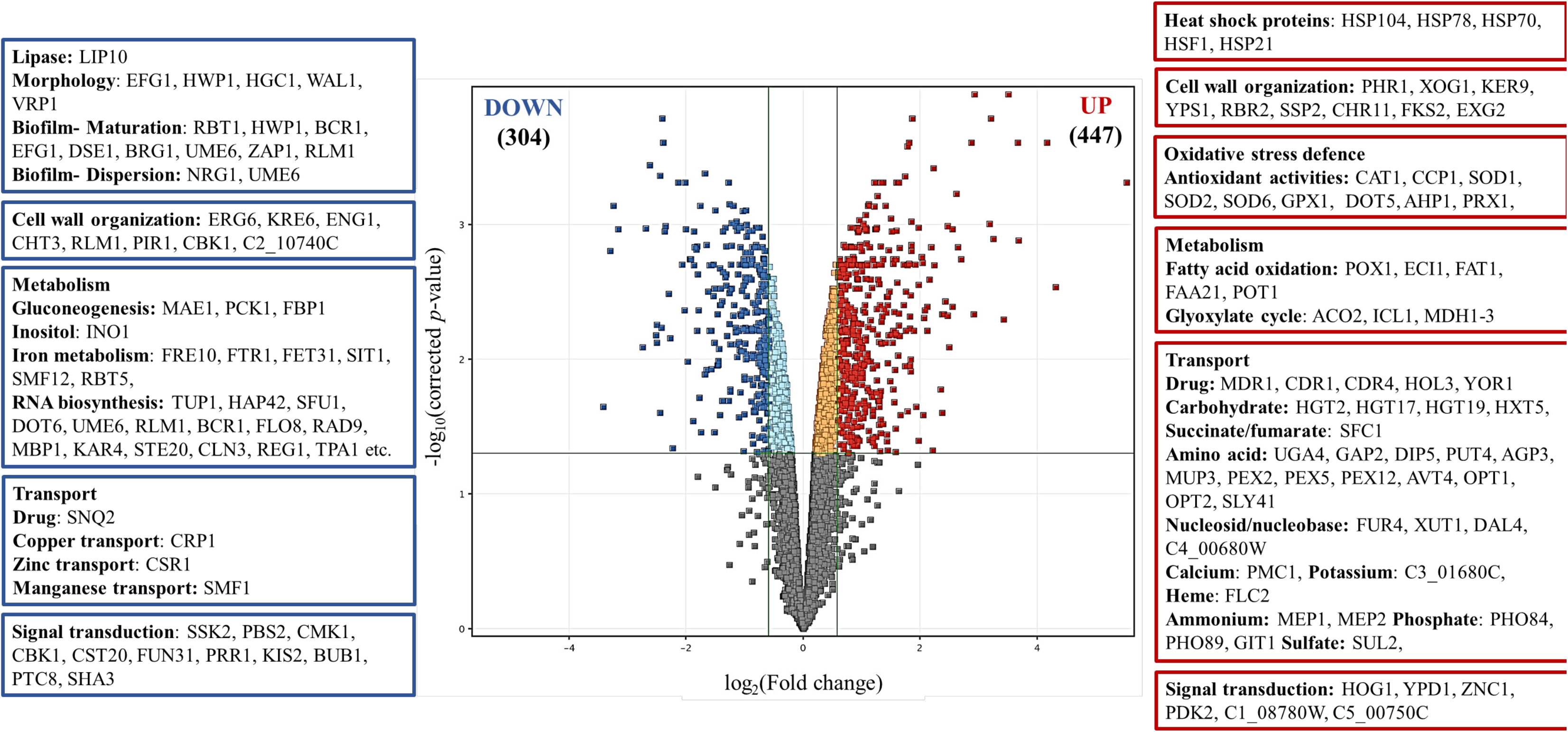
Overview of transcriptional changes induced by farnesol in *C. auris*. Upregulated (red) and downregulated (blue) genes were defined as differentially expressed genes (corrected *p* value of < 0.05), where more than 1.5-fold increase or decrease in their expression (farnesol treated *vs*. untreated). On the sides of the volcano plot are representative genes upregulated or downregulated by farnesol treatment. The data set is available in Supplementary Table 3.

**Figure 4.**
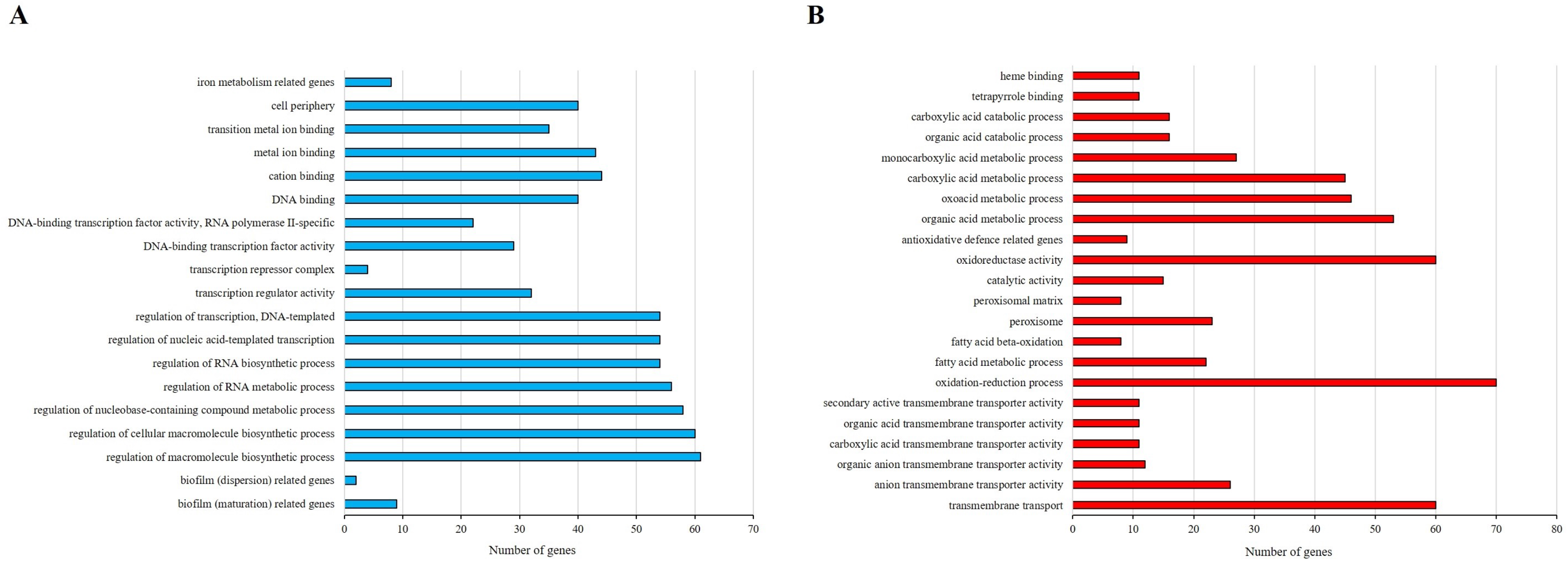
Summary of gene enrichment analyses and the number of genes affected by farnesol exposure of *C. auris.* Down-regulated (**A**, blue) and up-regulated (**B**, red) genes were defined as differentially expressed genes (corrected *p* value of <0.05). The enrichment of these gene groups were identified with the Candida Genome Database Gene Ontology Term Finder (https://www.candidagenome.org/cgi-bin/GO/goTermFinder) or were tested by Fisher’s exact test. The data set on the gene groups are available in Supplementary Tables 2 and 3.

### Evaluation of farnesol-responsive genes

To identify larger patterns in differential gene expression and to obtain an overall insight into the impact of farnesol, gene ontology terms were assigned to all of the genes in the *C. auris* genome; afterwards, we compared the terms for both the down-regulated and up-regulated genes to a background of all terms. We found 19 and 22 significant gene groups that were underrepresented and overrepresented in this analysis, respectively (Fig. 4, Tables S2 and S3).

#### Virulence-related genes

Virulence-related genes were significantly enriched within the farnesol-responsive down-regulated gene group, according to Fisher’s exact test (Table S3).

Most of these 11 putative genes are involved in biofilm maturation (*RBT1, HWP1, BCR1, EFG1, DSE1, BRG1, UME6, ZAP1,* and *RLM1*) and dispersion (*NRG1,* and *UME6*) (Figs. 3 and 4, Table S3); also, five down-regulated morphogenesis genes (*EFG1, HWP1, HGC1, WAL1,* and *VRP1*) are notable (Fig. 3, Table S3). Down-regulation of *RBT1* and *NRG1* under farnesol treatment was also supported by RT-qPCR data (Fig. 5 and Table S4).

**Figure 5.**
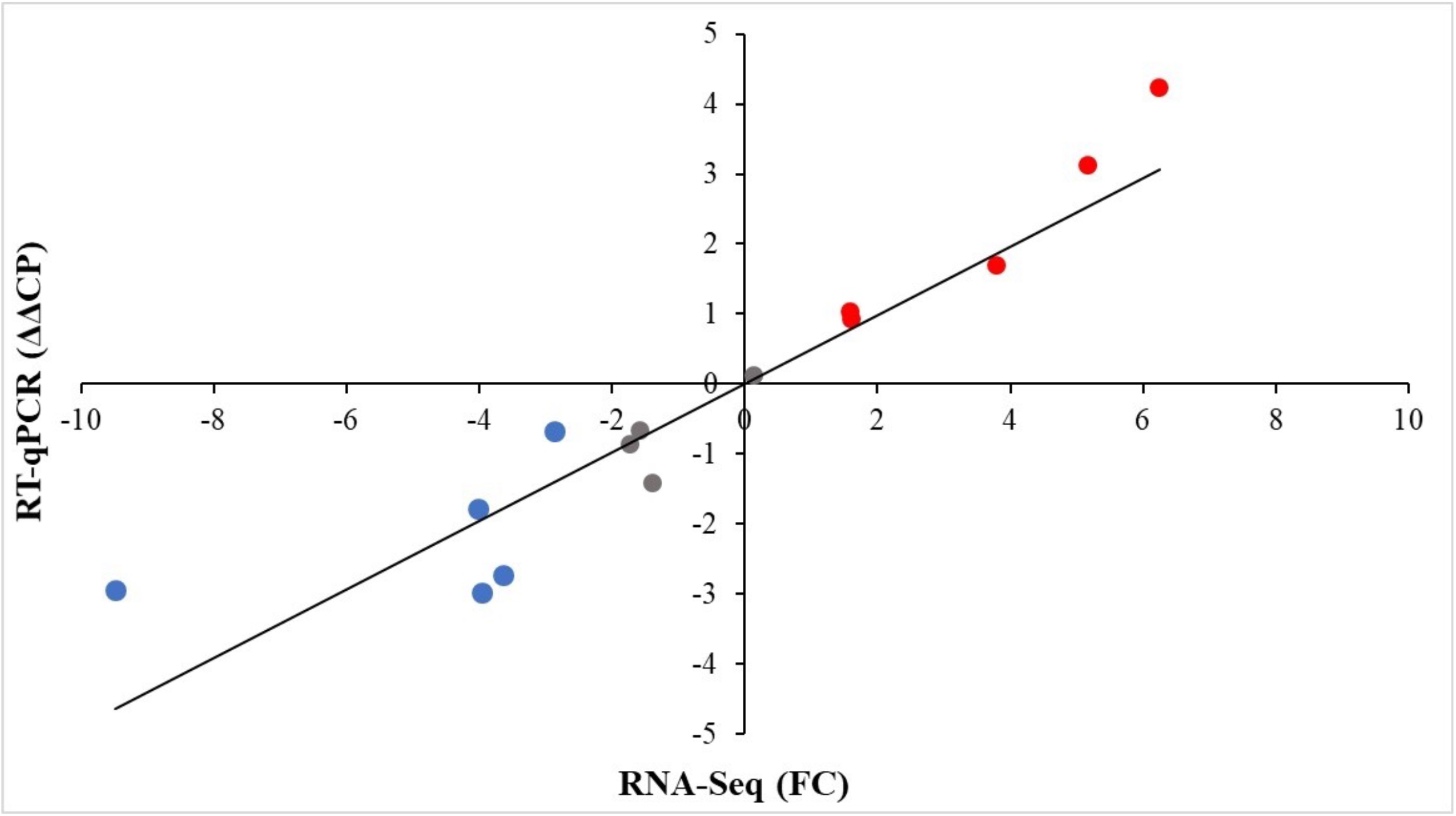
Correlation between RT-qPCR and transcriptome data. RNA-Seq data are presented as FC values, where FC is “fold change”. Relative transcription levels were quantified as ΔΔCP = ΔCP_control_ − ΔCP_treated_, where ΔCP_treated_ = CP_tested gene_ − CP_reference gene_, measured from farnesol treated cultures, and ΔCP_control_ = CP_tested gene_ − CP_reference gene_, measured from control cultures. CP values represent the qRT-PCR cycle numbers of crossing points. The *ACT1* gene was used as a reference gene. ΔΔCP values significantly (*p* < 0.05 by Student’s *t* test; n = 3) higher or lower than zero (up- or downregulated genes) are marked in red and blue, respectively. Pearson’s correlation coefficient between the RT-qPCR and RNA-Seq values was 0.87. The data set is available in Supplementary Table 4.

#### Oxidative stress-related genes

Genes belonging to antioxidative defense-related GO terms were enriched in the farnesol-responsive up-regulated gene group (Figs. 3 and 4, Table S3). Altogether, 9 genes were up-regulated after farnesol treatment, namely *CAT1, CCP1, SOD1, SOD2, SOD6, GPX1, DOT5, PRX1,* and *AHP1* (Fig. 3, Table S3). In addition, farnesol exposure increased the expression of *HSP21*, *YPD1,* and *HOG1*, encoding small heat shock protein, phosphorelay protein, and MAP kinase (Fig. 3, Table S3). Upregulation of *CAT1* and *CCP1*, coding for catalase and cytochrome-c peroxidase in farnesol-treated cells, was also confirmed by RT-qPCR (Fig. 5 and Table S4).

#### Metabolic pathway-related genes

Selected genes involved in glucose catabolism and fatty acid metabolism were determined with the *Candida* Genome Database (https://www.candidagenome.org). Farnesol treatment down-regulated *PCK1* and *FBP1*, encoding key enzymes specific to gluconeogenesis, but not glycolysis and tricarboxylic acid cycle genes (Fig. 3, Table S3). In addition, three genes related to the glyoxylate cycle (*ACO2*, *ICL1,* and *MDH1-3*) were significantly enriched in the up-regulated gene set (Fig. 3, Table S3).

Significant up-regulation of five putative genes encoding fatty acid β-oxidation enzymes was observed (*POX1, ECI1, FAT1, FAA21,* and *POT1*) (Figs. 3 and 4, Tables S2 and 3). In addition, farnesol exposure decreased the expression of *INO1*, encoding inositol-1-phosphate synthase (Fig. 3, Table S3).

Genes involved in iron homeostasis, including essential elements of reductive iron uptake (*FRE10, FET31, SMF12,* and *FTR1*), siderophore transport *(SIT1*), and hemoglobin use (*RBT5*), as well as manganese (*SMF1*, transporter), copper uptake (*CRP1,* transporter), and zinc metabolism (*CSR1*, transcription factor), were enriched in the down-regulated gene set (Figs. 3 and 4, Table S3).

The up-regulation of *POT1* (3-oxoacyl CoA thiolase) and the down-regulation of *INO1* and *FTR1* were supported by RT-qPCR data (Fig. 5 and Table S4).

#### Transmembrane transport-related genes

Farnesol treatment led to the increased expression of numerous genes (60 genes altogether) involved in transmembrane transport, including 5 putative antifungal drug transporter genes (*MDR1, CDR1, CDR4, HOL3,* and *YOR1*), 4 putative carbohydrate transport genes (*HGT2, HGT17, HGT19*, and *HXT5*), 13 putative amino acid transport genes, as well as 4 putative phosphate and sulfate transporter genes (*PHO84, PHO89, GIT1,* and *SUL2*) (Figs. 3 and 4, Table S3). Farnesol exposure also caused a significant increase in the expression of *CDR1* and *MDR1* (ABC transporters) as well as *HGT2* (glucose transmembrane transporter) of treated cells, according to the RT-qPCR results (Fig. 5 and Table S4).

### Effect of farnesol exposure on metal contents of *C. auris* cells

Farnesol treatment caused a significant reduction in intracellular iron, manganese, and zinc content (*p* < 0.05-0.001) compared to untreated control cells, whereas the level of intracellular copper was significantly increased, as shown in Table 1 (*p* < 0.001).

**Table 1.** Farnesol-induced iron, manganese, zinc and copper content in *Candida auris*.

| Metal contents/<br>Treatment | Mean value± SD <sup>a</sup> |  |  |  |  |
| --- | --- | --- | --- | --- | --- |
|  | Dry cell mass<br>DCM<br>(g/l) | Fe<br>(mg/kg) | Mn<br>(mg/kg) | Zn<br>(mg/kg) | Cu<br>(mg/kg) |
| Untreated<br>cultures | 0.38±0.08 | 152.2±21.1 | 67.9±5.1 | 787.8±22.2 | 274.6±15.7 |
| Farnesol-treated cultures | 0.09±0.01 *** | 116.0±10.0 * | 18.6±1.8 *** | 245.8±34.4 *** | 828.8±106.4 *** |

<sup>a</sup> Mean values ± standard deviations (SD) calculated from three independent experiments are presented.
\*and \*\*\* indicate significant differences at *p* values of <0.05 and 0.001, respectively, calculated by two-way ANOVA, comparing untreated control and farnesol-treated cultures.

## Discussion

Alternative treatments interfering with quorum-sensing have recently become attractive therapeutic strategies, particularly against difficult-to-treat multidrug-resistant pathogens such as *C. auris* (9,21–22). Previous studies have reported that fungal quorum-sensing molecules may have a remarkable antifungal effect and/or a potent adjuvant effect in combination with traditional antifungal agents (7,10,23–26). For example, Nagy et al. (2020) reported that supraphysiological farnesol exposure caused a significant reduction in the growth rate and metabolic activity of *C. auris* planktonic cells and biofilms, respectively (7). In addition, 75 μM farnesol treatment significantly decreased the fungal kidney burden in an immunocompromised systemic mouse model (7). Total transcriptome analysis using RNA-Seq may be an important technique to fully understand the underlying mechanisms of the observed antifungal effect exerted by these molecules.

Based on previous studies, farnesol induces a dose-dependent production of reactive species in *C. albicans*, especially at supraphysiological concentrations (27–28). In addition, these findings coincided with the *C. auris*-related physiological experiments published by Nagy et al. (2020) (7). In this study, several putative oxidative stress-responsive genes, namely *CAT1* (encoding for catalase activity), *GPX1* (encoding glutathione peroxidase), and *SOD1, SOD2, SOD6* (encoding superoxide dismutases), were up-regulated following exposure to farnesol. It is noteworthy that farnesol exposure also up-regulated *HOG1* MAP kinase, which is a critical component of the fungal oxidative stress response, further supporting the farnesol-induced oxidative stress in *C. auris* (29). This fact is further confirmed by the elevated DCF and superoxide dismutase levels in farnesol-exposed cultures (7).

Recent transcriptomic data have demonstrated that farnesol treatment affected the transcription of iron homeostasis-related genes, as well as the iron, zinc, manganese, and copper contents of *C. auris*. The down-regulation of iron uptake genes was associated with the significantly decreased iron content measured in farnesol-exposed cells. Similarly, the menadion sodium bisulphite induced oxidative stress also affected the transcription of iron homeostasis-related genes and the iron content of *C. albicans* cells (20). It should be noted that this response related to iron decrease may be a part of a general defense mechanism against farnesol and menadion sodium bisulphite to minimize the damage caused by ferrous ions. According to previous studies, elevated free intracellular iron levels facilitate the formation of reactive oxygen species and mediate iron-dependent cell death in *Saccharomyces cerevisiae* (20,30).

The down-regulated expression of *CSR1*, encoding a major transcription factor that stabilizes zinc homeostasis and provides cells with zinc-dependent protection against farnesol-induced oxidative stress (19), is related to the decreased intracellular zinc level observed. Zinc is an essential transition metal in oxidative stress defense because it is a structural component of superoxide dismutase, which is a key enzyme in the neutralization of superoxide radical anions (O_2_^●-^) (19).

In contrast to the majority of metals, manganese acts as an anti-oxidant element at high concentrations rather than a reactive oxygen species producer (19). However, farnesol inhibited the expression of *SMF1*, which is responsible for maintaining the intracellular manganese levels for anti-oxidant actions (19). In addition, the expression of *PMR1* (*p* < 0.05, FC = 1.2) was also inhibited, decreasing the virulence of fungal cells (31). This was associated with our previously published data, where daily farnesol treatment significantly decreased the virulence of *C. auris* (7).

The observed down-regulation of the copper exclusion system (*CRP1*/*CCC2*, encoding P‐type ATPases) may be associated with the significantly increased copper contents and the remarkable growth inhibition in farnesol-treated cells. Copper regulates a variety of cellular processes in fungal pathogens. When it presents in excess, it is associated with the generation of reactive oxygen species via the Fenton reaction and destroys the iron-sulfur cluster reducing the viability of cells (19,32–35). The elevated free copper levels in the farnesol-exposed cells may contribute to the increased redox imbalance quantified by DCF production (7), which was accompanied by increases in the specific activity of superoxide dismutase (7). Moreover, recent studies have shown that copper efflux pumps may be equally important in fungal defense strategies against phagocytes as for the virulence in *C. albicans* (19,32–35).

Interestingly, farnesol exposure exerted a significant up-regulation in several fatty acid β-oxidation-related genes (*POX1*, *ECI1*, *FAT1*, *FAA21*, and *POT1*). The elimination of unnecessary membrane lipids and the increased usage of fatty acids may provide a higher metabolic flux, needed for the maintenance of membrane fluidity (36). Jabra Rizk et al. (2006) and Rossignol et al. (2007) described that farnesol influences the membrane permeability in non-*albicans* species as *C. dubliniensis* and *C. parapsilosis* (5–6). The elevated fatty acid oxidation activity may explain the membrane-related farnesol effect, which may elucidate the previously observed antifungal effect (7). A further potential explanation of the antifungal effect can be found in the down-regulation of ergosterol biosynthesis-related genes, which alter the membrane permeability and/or fluidity (37). In our study, the *ERG6* gene was down-regulated following farnesol exposure, which may enhance the passive diffusion of farnesol across the membrane; furthermore, the decreased Erg6 content may confirm the higher susceptibility of *C. auris* cells to oxidative stress (37–38). Oliveira et al. (2020) showed that the *ERG6* mutant *Cryptococcus neoformans* displays impaired thermotolerance and increased susceptibility to oxidative stress as well as to different antifungal drugs, explaining, for instance, the previously reported synergizing effect with azoles (7). Furthermore, the *ERG6* mutant *C. neoformans* was totally avirulent in an invertebrate model, which may also explain the reduced virulence of *C. auris* after daily farnesol treatment (7,37). Beside *ERG6*, *INO1* was also down-regulated following farnesol treatment. This gene encodes the inositol-1-phosphate synthase, a key enzyme in the synthesis of inositol for phosphotidylinositol synthesis. The down-regulation of this gene may further explain the synergizing effect of farnesol with azoles against *C. auris* (7), because *INO1* is significantly up-regulated in drug-resistant *Candida* isolates (39).

With respect to the transport efflux pump-related genes, farnesol exposure caused a significant increase in the expression of *CDR1, CDR4, MDR1, HOL3,* and *YOR1*, whereas the expression level of *SNQ2* was decreased. Previous studies have revealed that these transporters mediate drug resistance for *C. auris* (40–41). Srivastava and Ahmad (2020) found that *CDR1*, *CDR2*, *MDR1*, *MDR2,* and *SNQ2* are significantly down-regulated in the presence of farnesol (42). Notably, there was a 1,000-fold difference between the farnesol dosages exerting the up-regulating effect (125 mM) compared to the concentration used in our study (75 μM). Nevertheless, our data support the hypothesis that farnesol, at lower concentrations, may be a potential substrate for the up-regulated transport proteins in order to protect the cells themselves from the oxidative stress induced by farnesol.

This is the first study analyzing the global changes in gene expression in *C. auris* following farnesol exposure, providing important insights into the mechanism of antifungal action of farnesol and the response of *C. auris*, facilitating a better understanding of farnesol-related antifungal activity. In summary, farnesol exposure enhanced the oxidative stress response and up-regulated drug efflux pumps, while reducing zinc and manganese intracellular content as well as iron metabolism. Moreover, cellular metabolism was modulated towards β-oxidation. These findings reveal the mechanisms underlying the antifungal effect and suggest that farnesol may represent a potent therapeutic option against this multi-resistant fungal superbug.

## Acknowledgements

R. Kovács was supported by OTKA Bridging Fund and FEMS Research and Training Grant (FEMS-GO-2019-502). R. Kovács was supported by the János Bolyai Research Scholarship of the Hungarian Academy of Sciences. F. Nagy was supported by the ÚNKP-19-3 and ÚNKP-20-3 New National Excellence Program of the Ministry for Innovation and Technology.

## Ethical approval

Not required

## Availability of data and materials

The RNA sequencing data discussed have been deposited in NCBI’s Gene Expression Omnibus (43) (GEO; https://www.ncbi.nlm.nih.gov/geo/) and are accessible through GEO Series accession number GSE 180093 (https://www.ncbi.nlm.nih.gov/geo/query/acc.cgi?acc=GSE180093).

---

**Supplementary Table 1**: Oligonucleotide primers used for RT-qPCR analysis.

**Supplementary Table 2**: Results of the gene set enrichment analysis.

Significant shared GO terms (*p* < 0.05) were determined with the Candida Genome Database Gene Ontology Term Finder (https://www.candidagenome.org/cgi-bin/GO/goTermFinder). Up- and down-regulated genes were defined as differentially expressed genes where log_2_(FC) > 0.585 or log_2_(FC) < −0.585. The FC ratios were calculated from the normalized gene expression values. Biological processes, molecular function and cellular component categories are provided.

**Supplementary Table 3**: Transcription data of selected gene groups.

Part 1: Genes involved in genetic control of *Candida auris* virulence.

Part 2: Genes involved in in selected metabolic pathways.

Part 3: Genes involved in ergosterol and fatty acid metabolism.

Part 4: Genes involved in response to oxidative stress.

Part 5: Genes involved in metals metabolism.

Part 6: Selected genes involved in regulation of RNA biosynthetic process.

Part 7: Selected genes have protein kinase or phosphatase activity.

Part 8: Selected genes involved in membrane transport.

The systematic names, gene names and the features (putative molecular function or biological process) of the genes are given according to the *Candida* Genome Database (https://www.candidagenome.org).

Up- and down-regulated gene were defined as differentially expressed genes with < 0.05 corrected *p* value. RNA-Seq data are presented as FC values, where FC is “fold change”.

Up- and down-regulated genes are marked with red and blue colour.

Results of gene enrichment analysis (Fisher’s exact test) are also enclosed to the parts 1-5. “The response of oxidative stress” genes (GOID: 0006979) collected from Gene Ontology Term Finder (https://www.candidagenome.org/cgi-bin/GO/goTermFinder).

**Supplementary Table 4**: Results of the RT-qPCR measurements.

Relative transcription levels were quantified with ΔΔCP = ΔCP_control_ – ΔCP_treated,_ where ΔCP_treated_ = CP_tested gene_ – CP_reference gene_ was measured from treated cultures and ΔCP_control_ = CP_tested gene_ – CP_reference gene was_ measured from control cultures. CP values represent the qRT-PCR cycle numbers of crossing points. RT-qPCR data are presented as mean ± SD calculated from three independent measurements, normalised to the *ACT1* gene expression and were compared using Student’s *t*-test (*p*<0.05). Significantly higher or lower than zero ΔΔCP values (up- or down-regulated gene) are marked with red and blue colours.

